# Habitat heterogeneity, environmental feedbacks, and species coexistence across timescales

**DOI:** 10.1101/2022.06.26.497662

**Authors:** Zachary R. Miller, Stefano Allesina

**Affiliations:** Department of Ecology & Evolution, University of Chicago, Chicago, IL, USA; Northwestern Institute on Complex Systems, Evanston, IL, USA

## Abstract

A large body of ecological theory explains the coexistence of multiple species in variable environments. While spatial variation is often treated as an intrinsic feature of a landscape, it may be shaped and even generated by the resident community. All species modify their local environment to some extent, driving changes that can feed back to affect the composition and coexistence of the community, potentially over timescales very different from population dynamics. We introduce a simple, nested modeling framework that describes species coexistence in heterogeneous environments, as well as the evolution of this heterogeneity over time due to feedbacks from the biotic community. We derive analytical conditions for the coexistence of any number of species in environments with intrinsic heterogeneity or feedbacks, and identify essential differences between these scenarios. Our model is naturally simplified in the limit of very fast or very slow environmental feedbacks, allowing us to treat these two scenarios – which bookend the full range of dynamics – in detail. Among other results, we demonstrate how dispersal and environmental specialization interact to shape realized patterns of habitat association. We also show that environmental feedbacks can tune landscape conditions to promote stable coexistence, although feedbacks can give rise to complex dynamics such as limit cycles, priority effects, and episodic dynamics, as well. Our flexible modeling framework helps explain how and when each of these behaviors arise, and offers a generic mathematical platform for exploring the interplay between species and landscape diversity.

## 1 Introduction

nvironmental heterogeneity provides the raw material for niche partitioning in ecological communities. When the environment varies from place to place, differences in the ways species respond to local conditions can facilitate their coexistence at the landscape scale, even when local coexistence is impossible (Chesson 2000; Amarasekare 2003). This connection between environmental heterogeneity and the maintenance of species diversity has deep roots in ecology (Andrewartha *et al*. 1954; MacArthur *et al*. 1966; Whittaker 1967), and has been well-studied both theoretically (Horn & MacArthur 1972; Chesson 1985; Iwasa & Roughgarden 1986; Chesson 2000) and empirically (Schoener 1974; Silvertown *et al*. 1999; Codeco & Grover 2001; Oliver *et al*. 2010).

However, this picture becomes more complex when species themselves shape their environment. Feedbacks between the biotic community and landscape conditions across space can enhance or reduce environmental heterogeneity over time (Crooks 2002; Wright *et al*. 2002; Pastor 2005). Prominent examples include plant-soil feedbacks (Bever *et al*. 1997; Mangan *et al*. 2010; Van der Putten *et al*. 2013) and related Janzen-Connell effects Janzen (1970); Connell (1971), wherein plants directly or indirectly shape the local densities of soil microbes or natural enemies such as seed predators, generating a dynamic landscape of legacy effects. These processes are thought to play an important role in maintaining the diversity of many natural plant communities, but they may also lead to positive feedbacks that drive monodominance Wolfe & Klironomos (2005); Van der Putten *et al*. (2013).

Here, we introduce a flexible modeling framework for community dynamics in heterogeneous landscapes with and without feedbacks that change environmental conditions through time. We build on the classical metapopulation paradigm Levins (1969) and related patch models, which provide a minimalist approach to studying ecosystems with distinct local and global scales Chesson (1985); Iwasa & Roughgarden (1986); Amarasekare (2003). The simplicity of this framework allows us to capture and analyze essential features of complex environmental feedbacks. While a range of conceptual and quantitative models (Roughgarden 1978; Bever *et al*. 1997; Gurney & Lawton 1996; Hui *et al*. 2004; Raynaud *et al*. 2013; Monk & Schmitz 2021) have shed light on such processes – and particularly on when they might help or hinder coexistence – they remain challenging to incorporate in tractable mathematical frameworks. One central obstacle is the high species diversity in many natural systems, although accounting for this diversity is crucial to understanding real-world coexistence (Miller *et al*. 2021). Additionally, many studies – and even sub-fields of ecology – focus on a single source of heterogeneity, making it hard to draw general conclusions that cut across system specifics. In particular, fixed or “exogenous” heterogeneity and biotically-generated or “endogenous” heterogeneity have often been approached from very different perspectives (Bolker 2003; Smith 2022). Our models and analysis help overcome these challenges by providing coexistence criteria that extend naturally to communities of arbitrary size, and by building endogenous environmental feedbacks directly into a core model for landscapes with exogenous heterogeneity.

We develop a general approach that is agnostic to the specific sources of environmental heterogeneity and allows us to consider feedbacks on a wide range of timescales. We show that our framework includes a recent model for rapid habitat modification (Miller & Allesina 2021) as a limiting case. To delineate the full spectrum of possible dynamics, we focus especially on the the opposite extreme of feedbacks that shape the landscape over very long times. In doing so, we identify essential features of coexistence maintained by exogenous and endogenous heterogeneity. Across the two extremes of very fast and very slow feedbacks, coexistence maintained by endogenous heterogeneity can be characterized by the same simple analytical conditions. Over long times, these limiting cases behave similarly to one another, and qualitatively differently from systems with exogenous heterogeneity, even though slow feedbacks are difficult to distinguish from exogenous heterogeneity on short timescales. Our results help explain how environmental feedbacks emerge and play out over different timescales, and the modeling framework we introduce offers a platform for comparing exogenous and endogenous heterogeneity on equal footing. Finally, we discuss how and under what conditions our models can be parameterized from observational data, and other implications for the analysis of community dynamics in natural communities.

## 2 Exogenous heterogeneity

We consider a landscape composed of many local patches, which can be classified into *ℓ* discrete types. The type of a patch summarizes its internal conditions as they are relevant to a focal community of *n* species inhabiting the landscape. For example, in the context of a plant community, patches might be classified by soil type or topography. The rate at which each species can establish in a patch depends on the patch type, and may differ between species, reflecting interspecific differences in niche requirements. For simplicity, we follow conventional models and assume there is global dispersal between all patches, and that each patch can be occupied by at most a single species at any time. Denoting the proportion of all patches that are of type *j* and occupied by species *i* at time *t* by *X*_*ij*_(*t*), and the proportion of all patches that are of type *j* and vacant by *y*_*j*_(*t*), we can model the dynamics of these proportions across a sufficiently large landscape by:

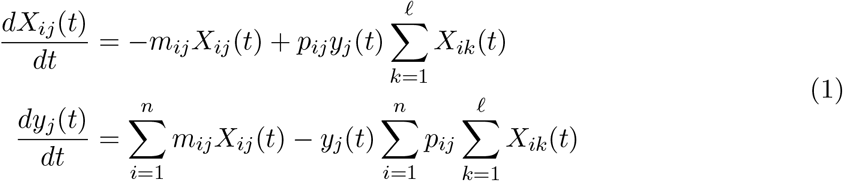

The parameters *p*_*ij*_ ≥ 0 and *m*_*ij*_ *>* 0 specify the rates at which species *i* establishes or goes locally extinct in patches of type *j*, respectively. As in classic metapopulation models, the dynamics represent the net action of these two processes: colonization of empty patches by propagules or dispersers from occupied patches, and local extinctions at a constant rate per patch. The summations over patch type (index *k*) reflect the fact that empty patches of type *j* may be colonized by propagules of species *i* dispersed from patches of any type. Summations over species identity (index *i*) reflect the fact that the type is a fixed property of each patch, so a patch of type *j* always returns to the *y*_*j*_ class when vacated. This also implies that the total proportion of patches of type *j*, which we denote by *w*_*j*_ = *y*_*j*_(*t*) + ∑_*i*_ *X*_*ij*_(*t*), is constant through time for every *j*.

In principle, we can allow local extinction rates to depend on both species identity and patch type; however, in this study we focus primarily on the simplest case where *m*_*ij*_ = *m*. Thus, the effects of landscape heterogeneity are realized through rates of establishment, not local persistence. The suitability of this assumption will depend on the community of interest, but environmental heterogeneity is thought to act more strongly on establishment than persistence in many systems, typically because smaller populations or immature individuals are more sensitive to patch conditions (Grubb 1977; Kryazhimskiy *et al*. 2007; Mächler & Altermatt 2012; Baldeck *et al*. 2013). We are also assuming that there are no significant differences in local extinction rates between species, reflecting a community of demographically similar species, or a system where local extinctions are primarily driven by external disturbance.

Under this equal *m* assumption, we can greatly simplify the dynamics by tracking only the proportion of patches – regardless of type – occupied by species *i* at time *t*. We denote these proportions by *x*_*i*_ =∑ _*j*_ *X*_*ij*_. Instead of *ℓ* × (*n* + 1) equations, there are now *n* + *ℓ*, given by:

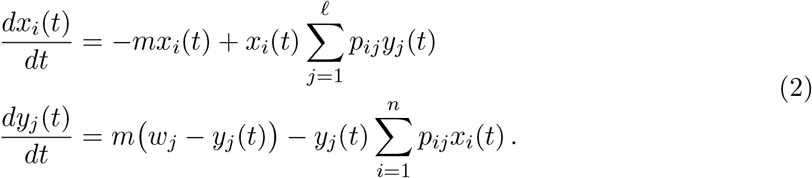

The *w*_*j*_, which must sum to one, are now seen as parameters that describe the heterogeneity of the landscape.

These dynamics take precisely the same mathematical form as a consumer-resource model with external inflow of abiotic resources (Tilman 1980; Butler & ODwyer 2018; Marsland III *et al*. 2020). In this parallel, each species plays the role of a consumer, and each patch type is interpreted as a resource with inflow proportional to *w*_*j*_. Both “consumers” and “resources” experience density-independent mortality at a rate *m*, and the *p*_*ij*_ are analogous to consumption rates.

Consumer-resource systems of the form in Eq. 2 have been studied extensively, allowing us to immediately draw conclusions about multispecies dynamics in heterogeneous metapopulations by translating results from the consumer-resource setting. For example, it is well-known that the number of coexisting consumers is at most equal to the number of resources in such models, a classic result known as the competitive exclusion principle (Levin 1970). Our model equivalence provides a formal demonstration that this intuitive principle carries over to the context of environmental heterogeneity. Given this upper limit, we study coexistence assuming *ℓ* = *n*, representing a fully “packed” consumer community. In this case, Eq. 2 has at most a single *coexistence equilibrium*, which is easily expressed in matrix form:

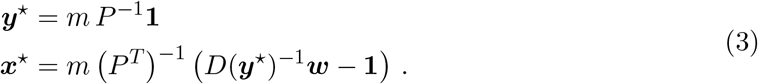

Here we use the notation ***y***^⋆^ for the vector of equilibrium proportions (and similarly for ***x***^⋆^ and ***w***), and we collect the coefficients *p*_*ij*_ in the matrix *P*. We also use *D*(***y***^⋆^) for the diagonal matrix with non-zero elements given by ***y***^⋆^, and **1** for a vector of *n* ones.

Many properties of this equilibrium are known. Coexistence of all *n* species requires that the equilibrium is biologically feasible, meaning that the components of ***y***^⋆^ and ***x***^⋆^ are all positive. In fact, it can be proven that this equilibrium is globally stable whenever it is feasible, and therefore coexistence is entirely controlled by feasibility (see, e.g., Marsland III *et al*. (2020); for completeness, we provide a proof of this result in SI Appendix 1.3).

In accordance with the fact that ***x*** and ***y*** are proportions in our original framing of the model, we find that feasibility of ***x***^⋆^ requires 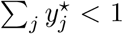. This means that there is an upper limit on *m* imposed by the matrix of establishment rates: *m <* **1**^*T*^ (*P* ^*−*1^)**1**. Remarkably, the vacant patch equilibrium values, ***y***^⋆^, are completely unaffected by the underlying distribution of patch types in the environment, ***w***. As in consumer-resource models, this “shielded” behavior arises because the species in the system robustly drive the supply of vacant patches to a point determined only by their own demographic rates (Tikhonov & Monasson 2017). Feasibility of these values requires that every element of *P* ^*−*1^**1** is positive. Loosely, in ecological terms this expresses a requirement that species are sufficiently similar in overall colonization ability (i.e., *p*_*ij*_ averaged across patch types), or sufficiently specialized on distinct habitat types (see SI Appendix 1.2 for detailed interpretation of this condition).

The equilibrium proportions for species themselves *are* sensitive to ***w***, but indirectly. Even if each species is specialized on a different patch type, the ***x***^⋆^ and ***w*** are not simply proportional, unless the species are all perfect specialists (Tikhonov & Monasson 2017). For example, in Fig. 2, we illustrate a scenario where each species is sufficiently specialized so that all non-preferred patches are net sinks (i.e., *p*_*ij*_ *< m* for all *i* ≠ *j*). Changing the distribution of patch types in the landscape can cause some species to go extinct, even though patches of all types remain available. Counterintuitively, this means that increasing the diversity (evenness) of habitat types will often, and sometimes drastically, decrease the species diversity of the system. This kind of unimodal relationship between habitat heterogeneity and diversity is often observed in natural ecosystems Ben-Hur & Kadmon (2020), and our model illustrates a simple underlying mechanism. Strong competitive interference arises in our model, and can constrain species coexistence, because the joint distribution of species and patch types reflects not just habitat preference, but also source-sink dynamics between patches. The potential for such dynamics in heterogeneous landscapes has long been noted (Horn & MacArthur 1972; Holt 1997; Shurin *et al*. 2004).

**Figure 1:**
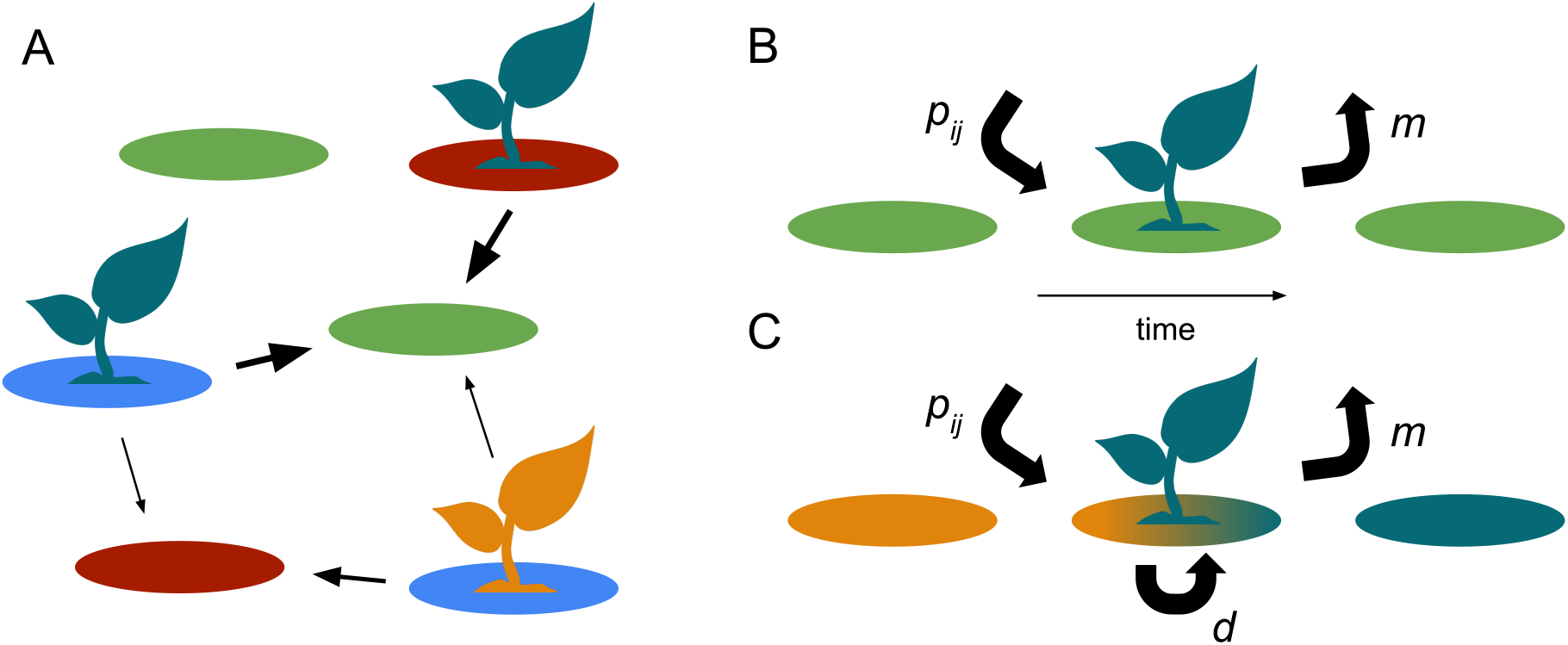
Metapopulation models with exogenous and endogenous patch heterogeneity. (A) We model an ecosystem where species disperse between patches with varying local conditions. The environmental conditions within a patch, summarized by the patch type or state, influence the rates at which different species can colonize and establish. We consider models where variation in patch conditions is (B) a fixed property of the landscape (Eq. 2), or (C) shaped by the biotic community over time (Eq. 5). Here, colors indicate patch types/states and species identities. When heterogeneity is endogenous (C) each patch state is identified with a species in the community, reflecting environmental modifications (occurring at a rate *d*) due to that species’ presence.

**Figure 2:**
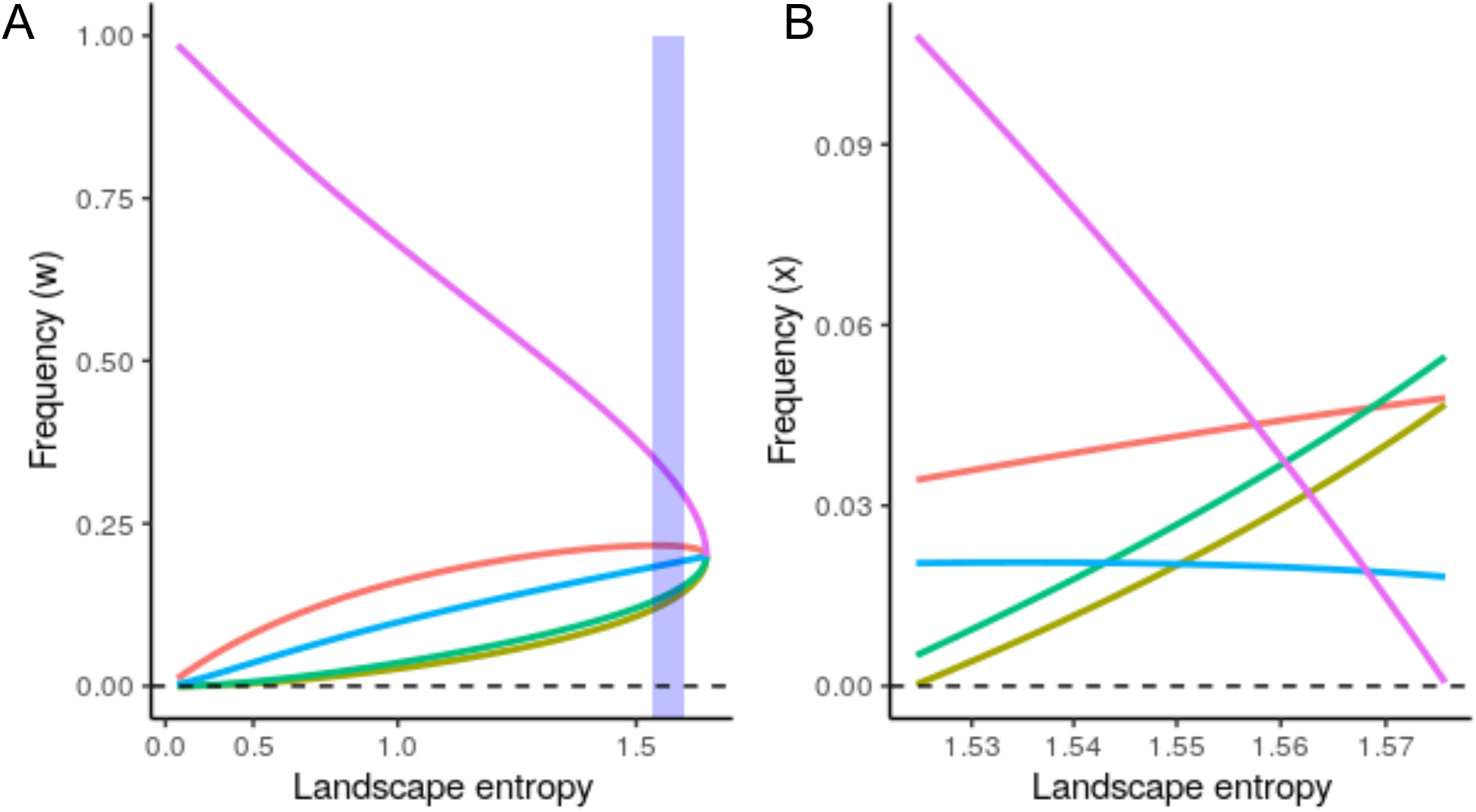
We illustrate the relationship between landscape parameters (A) and species’ equilibrium frequencies (B) using a simulated community of habitat specialists. We sampled *P* such that each species had a strong advantage colonizing a distinct patch type, and *p*_*ij*_ *< m* for non-preferred patches (parameters available at (Miller & Allesina 2022)). We then calculated species’ frequencies (***x***^⋆^) for landscapes (***w***) ranging from a uniform environment dominated by one patch type, to equal frequency of all types. The full gradient of ***w*** parameters is shown in (A); the five species community is only feasible for a small subset (blue shading). In (B), we show species’ frequencies, as a function of landscape diversity (measured in Shannon entropy), for the feasible range. Species’ frequencies are not straightforwardly related to the availability of their preferred habitat types (indicated by matching colors), and species go extinct as ***w*** varies, even though all patch types are present. Surprisingly, even increasing the diversity (evenness) of the landscape drives some species extinct. To generate a gradient of ***w***, we selected parameters ***v*** compatible with feasibility and let 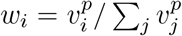 varying *p* from 0 to 10.

In our model, we can examine this joint distribution more closely by returning to the “full” dynamics in Eq. 1. The hierarchical structure of Eqs. 1 and 2 immediately implies that each *X*_*ij*_ reaches a stable equilibrium value

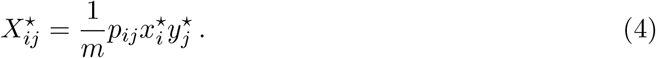

Here, and in Fig. 3, we see clearly that the net pattern of patch occupancy in an equilibrial landscape is shaped by the particular rates at which a focal species *i* can colonize and establish in patches of type *j* (*p*_*ij*_), as well as the overall abundance of *i* and availability of *j* in the system 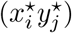. These two factors can be identified with “species sorting” and “mass effects” processes, respectively, which are usually viewed as two ends of a continuum for metacommunity dynamics (Leibold *et al*. 2004; Shoemaker & Melbourne 2016). While many studies of metacommunities focus on determining the importance of one or the other process in a particular community, our model quantitatively predicts how the two should act together to shape observed patterns of habitat association.

**Figure 3:**
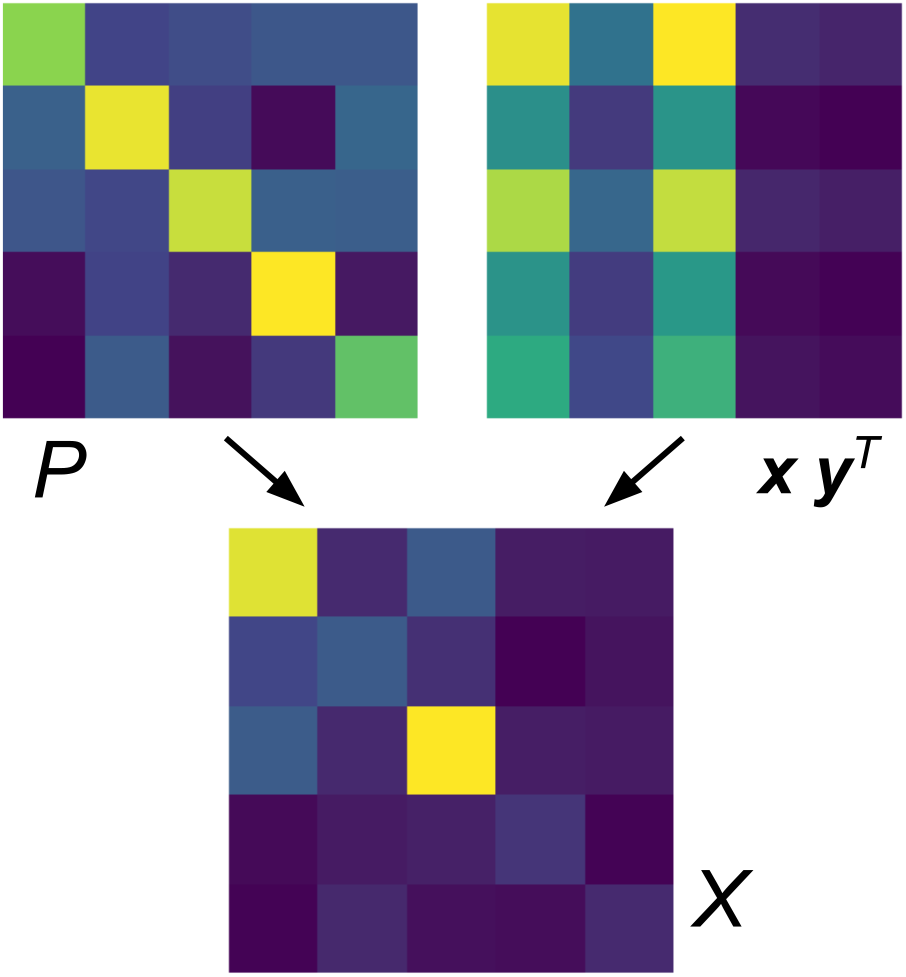
Pattern of habitat association reflects combination of species sorting and mass effects. The joint distribution of species and patch types at equilibrium, 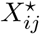 (Eq. 4), at bottom, is shaped by the matrix of colonization and establishment rates *P* (left) and the overall abundance of species and vacant patch types in the ecosystem ***xy***^*T*^ (right). The left pattern sets an expectation based on species’ habitat preferences (in this example, strong specialization on distinct patch types), while the right pattern shows the expected distribution if patches were homogeneous. The actual pattern of habitat association is the element-wise product of the two. Here each matrix is normalized so that the elements sum to one. Higher values are indicated by light colors (yellow), and low values by dark colors (blue).

## 3 Endogenous heterogeneity

Next, we consider how community dynamics change when patch heterogeneity is no longer a static feature of the environment, but an outcome of biotic feedbacks. For simplicity and for compatibility with the framework developed for exogeneous heterogeneity, we maintain the assumption that patches can be classified into distinct types, although now we speak of patch *states*, since these attributes change through time. Each patch state is associated with a distinct species, reflecting the impacts of that species on local environmental conditions. For example, we might consider distinct soil microbial communities associated with particular host plants (Bever 2002; Schweitzer *et al*. 2008; Philippot *et al*. 2013), immune statuses corresponding to recent infection history in vertebrate hosts (Kucharski *et al*. 2016), substrate morphologies shaped by benthic “ecosystem engineers” (Levinton 1995; Gutiérrez *et al*. 2003), or chemical concentrations maintained by different bacterial strains (Ratzke & Gore 2018; Amor *et al*. 2020). Thus, the number of species and patch states are both *n*, and we assume these are labelled so that state *i* corresponds to legacy effects of species *i*. As before, patch states affect the dynamics of the community by governing the rate of colonization and establishment by each species.

We approximate changes in local environmental conditions as discrete shifts between patch states. In a patch occupied for some time by species *i*, we assume that the local population of species *i* can drive a transition from the current patch state to state *i* at some rate *d*. In principle, this rate might depend on both the current patch type and the identity of the occupant species, but we focus on the case where *d* is constant. This scenario naturally describes systems where some external disturbance, occurring at an approximately constant rate across the landscape, is needed to shift patches between alternative states, or where species all species modify the environment at roughly equal rates.

With biotic feedbacks operating, the distribution of patch states in the landscape, ***w***, is now a dynamic variable. But given a particular distribution of patch states ***w***(*t*) at time *t*, we assume the instantaneous dynamics of colonization and extinction are exactly the same as in Eq. 1. This leads to the following model for a community with endogeneous heterogeneity:

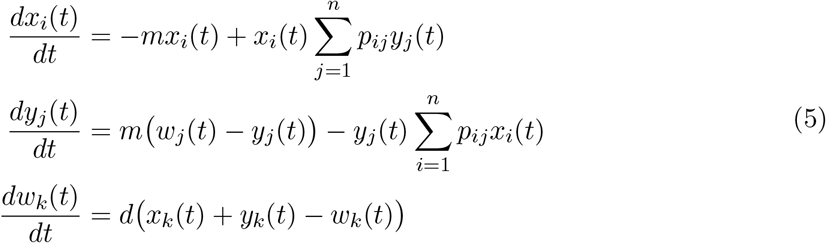

Intuitively, the total proportion of patches in state *k* increases when the number of patches occupied by species *k*, regardless of state, exceeds the total number of occupied patches in state *k*, regardless of occupant (given by *w*_*k*_(*t*) − *y*_*k*_(*t*)). A detailed derivation of Eq. 5) is given in SI Appendix 2.1.

The dynamics of Eq. 5 can be much more complex than Eq. 2 – potentially including non-equilibrium coexistence or multistability – making it difficult to fully characterize the behavior of this model. To make progress, we first consider two limiting cases where patch dynamics and underlying landscape dynamics operate on very different timescales, permitting a natural simplification of the model via fast-slow decomposition (Tikhonov 1952; ODwyer 2018).

In the first case, all species modify their local environment extremely rapidly. Consequently, the state of a vacant patch will invariably reflect the identity of the most recent resident species. Formally, this scenario represents the limit where *d* → ∞, and we can apply a fast-slow decomposition. Treating ***x***(*t*) and ***y***(*t*) as fixed, because these variables change slowly compared to patch states, ***w***(*t*), the dynamics 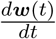 become a set of *n* decoupled differential equations, each with a stable equilibrium where *w*_*k*_(*x*_*k*_(*t*), *y*_*k*_(*t*)) = *x*_*k*_(*t*) + *y*_*k*_(*t*). We take ***w*** to be at this equilibrium at all times (i.e. we let 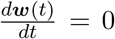 and substitute in the dynamics for ***y***(*t*) to obtain the *slow system*:

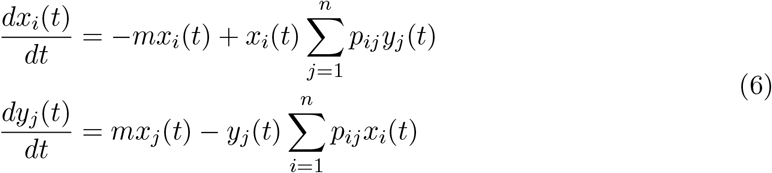

In fact, this system is precisely the habitat modification model we have studied previously (Miller & Allesina 2021).

At the opposite extreme, patches change state very slowly (that is, rarely). This limit corresponds to a system where patches resist modification – for example, where patch states are alternative stable states (Petraitis & Latham 1999; Amor *et al*. 2020) – but species occasionally drive patches from one state to another. Again, we consider a fast-slow decomposition, now with *d* → 0. Taking ***w***(*t*) as fixed over short timescales, we obtain a *fast system* that is identical to Eq. 2. Provided the equilibrium (Eq. 3) for this system is feasible, we have seen that it is globally stable, so we take ***x***(*t*) and ***y***(*t*) to be at this equilibrium at all times. Then we find the slow system for the gradual evolution of patch states:

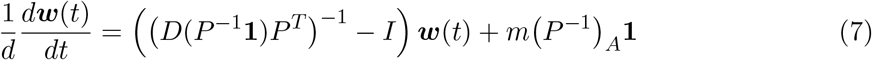

Here *I* is the identity matrix of size *n* and *S*_*A*_ denotes the anti-symmetric part of a matrix, *S*_*A*_ = *S* − *S*^*T*^. This system is a matrix differential equation, which is very amenable to analysis. In SI Appendix 2.2, we prove that Eq. 7 has a unique equilibrium solution, ***w***^⋆^, which is always feasible if ***y***^⋆^ = *mP* ^*−*1^**1** is feasible, and which guarantees the feasibility of ***x***^⋆^. This equilibrium may be stable or unstable, depending on the structure of the colonization rate matrix *P*. We derive a simple graphical condition that characterizes the stability of Eq. 7 in terms of the eigenvalues of *PD*(*P* ^*−*1^**1**), which is a weighted version of *P* reflecting the distribution of vacant patch states at equilibrium (see Fig. S1). This normalization by *P* ^*−*1^**1** removes any effect of the overall quality of different habitat types.

In the special case where *P* is symmetric (i.e., *p*_*ij*_ = *p*_*ji*_), the analysis and interpretation of the model become even simpler. This kind of symmetry may arise naturally (at least approximately) in systems where the effect of one species’ habitat modifications on another species’ establishment rate depends on some measure of similarity between them. For example, the degree of “spillover” of Janzen-Connell effects between two tropical tree species is a function of their phylogenetic relatedness (Gilbert & Webb 2007). In this symmetric case, ***x***^⋆^, ***y***^⋆^, and ***w***^⋆^ all become proportional to *P* ^*−*1^**1**. This equilibrium is stable if and only if *P* has exactly one positive eigenvalue. We prove this result in SI Appendix 2.2 and also show that this characterization of stability extends somewhat beyond the context of strict symmetry: For arbitrary *P*, if exactly one eigenvalue of *PD*(*P* ^*−*1^**1**) lies in the right half of the complex plane, then the coexistence equilibrium will be stable (although this condition is only *necessary* for stability if *P* is symmetric).

Precisely the same stability condition characterizes the dynamics of Eq. 6 when *P* is symmetric Miller & Allesina (2021). The appearance of this stability criterion at both limiting extremes of Eq. 5 suggests that it applies more broadly to feedbacks on any timescale. Indeed, in SI Appendix 2.3, we prove that this condition characterizes local stability of the coexistence equilibrium when *P* is symmetric for *any* value of *d*. This stability condition can be seen as a quantitative generalization of the notion that each species must modify the environment in a way that disadvantages itself in order to generate negative frequency-dependent feedbacks that maintain diversity. This interpretation is grounded in two mathematical facts: First, for this eigenvalue condition to hold, it is necessary that *p*_*ij*_ ≥ min(*p*_*ii*_, *p*_*jj*_) for all *i* and *j*. Second, if 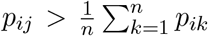 for all *i* ≠*j*, then the condition is guaranteed to hold. These necessary and sufficient characterizations both effectively limit the rate at which species can colonize patches modified by conspecifics, in agreement with classic conceptual arguments Janzen (1970); Connell (1971); Bever *et al*. (1997).

Aside from this intuitive stability condition, there are other qualitative similarities between the dynamics with very fast and very slow feedbacks. In particular, we have shown that feasibility in both cases depends only on ***y***^⋆^; there is always a distribution of patch states ***w***^⋆^ compatible with feasibility, and the system will evolve toward this configuration when a stability criterion is met. This behavior stands in contrast to the model with exogenous heterogeneity (Eq. 2), where any feasible equilibrium is stable, but feasiblity depends strongly on ***w***.

The dynamics of fast and slow feedbacks are not entirely identical, however. The fast- feedback dynamics (Eq. 6) can exhibit stable limit cycles when *P* is not symmetric Miller & Allesina (2021). Because Eq. 7 is a linear system, this behavior is impossible when feedbacks occur on very long timescales. But as *d* → 0, the model can still exhibit complex dynamics in the form of long transients (Hastings *et al*. 2018). In Fig. 5, we illustrate two interesting and ecologically relevant behaviors that can arise. First, if *P* meets the stability condition for the slow system (Eq. 7), then the dynamics will eventually reach a stable, feasible equilibrium. However, ***w***(*t*) may transiently take on values that are incompatible with the feasibility of some species (i.e., some *x*_*i*_(*w*(*t*)) *<* 0 for a range of *t*). In this case (Fig. 5A), the dynamics will “jump” between feasible subsystems where some species reach extremely low frequencies. In natural systems with finite size, this might lead to the extinction of these species before the coexistence equilibrium is reached. Alternatively, in very large systems or where there is a source of immigration that can rescue populations from rarity, the dynamics will be highly episodic as the system abruptly switches between feasible states, potentially over long times.

**Figure 4:**
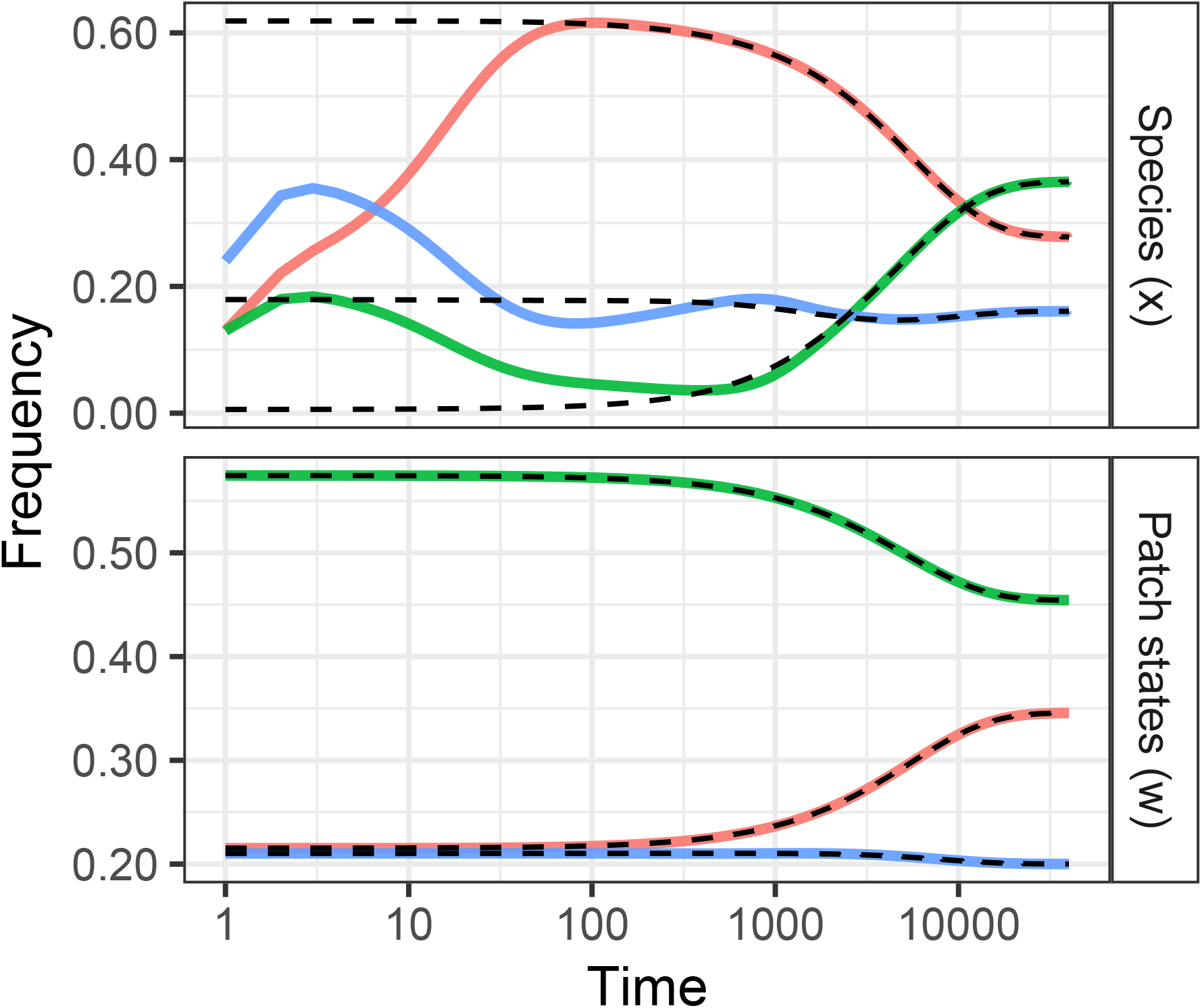
Stable feedback dynamics converge to fast-slow limit. Here we show the evolution of species frequencies (top) and the distribution of patch states (bottom) over time when feed-backs are slow. Colored lines indicate the solution obtained by numerically integrating Eq. 5 with *d* = 10^*−*5^. Dashed lines show the predicted dynamics using the fast-slow decomposition. The predictions for patch states were obtained by solving the slow system (Eq. 7), and the predictions for species’ frequencies were calculated from the predicted ***w***(*t*) using Eq. 3 at each time point. The dynamics rapidly collapse to the slow manifold derived analytically. In these simulations, *m* = 0.1 and the elements of *P* were sampled symmetrically *iid* from the standard uniform distribution until a feasible and stable parameter set was found (parameters available at (Miller & Allesina 2022)). Note that time is shown on a log scale, to highlight dynamics on long and short timescales.

**Figure 5:**
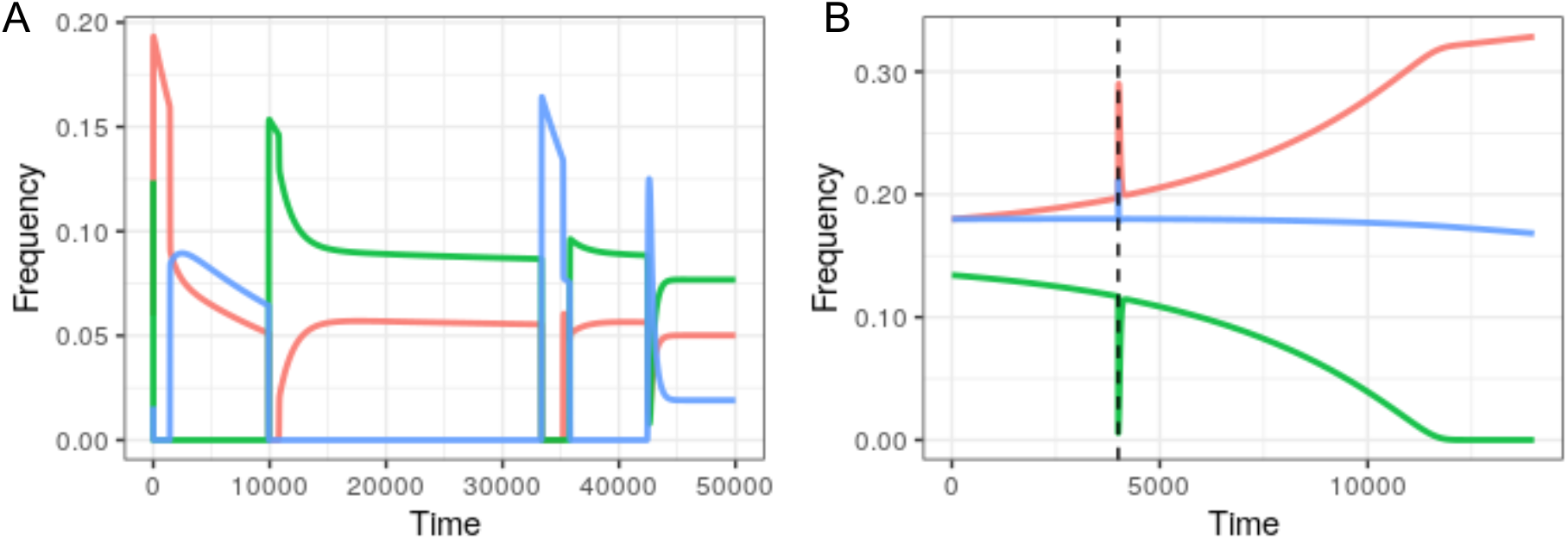
Complex transient dynamics when environmental feedbacks are much slower than community dynamics. Here we show species’ frequencies (***x***(*t*)) for the two scenarios described in the text. (A) When the slow system (Eq. 7) is stable, the ecosystem will eventually evolve toward a distribution of patch states such that species coexistence is feasible. However, if ***w*** is transiently incompatible with feasibility of the full community, the system will jump between (transiently) feasible sub-communities, leading to highly episodic dynamics. Each species reaches very low abundances during the transient dynamics, which would lead to extinctions in the absence of immigration or storage mechanisms (e.g., seed banks). (B) When the slow system is unstable, one or more species will eventually be excluded, but very gradually. During the transient period, the frequencies of all species are robust to perturbations. Here, the green species, which is driven to extinction by the long-term feedback dynamics, rapidly recovers from reduction to near extinction (dashed line) early in the dynamics. In these simulations, *m* = 0.5, *d* = 10^*−*5^, and the elements of *P* were sampled *iid* from the standard uniform distribution until the desired properties were found (parameters available at (Miller & Allesina 2022)).

If instead *P* is not compatible with stability, some species will eventually be excluded from the system due to long-term feedbacks between the community and the landscape. On short timescales, however, the distribution of patch states is approximately constant, and the dynamics of Eq. 5 will closely resemble Eq. 2. In particular, species frequencies will be stable to perturbations. Thus, we can find surprising scenarios where a species rapidly recovers from reduction to low abundance in the short term, even though it is ultimately doomed to a gradual extinction (Fig. 5B).

These two scenarios illustrate that species’ ability to invade when rare can change at different points in the dynamics, as the community modifies conditions across the landscape.

## 4 Discussion

Environmental heterogeneity is commonly understood to beget species diversity, but the converse can also be true. To better understand the relationships between the two, we introduced a simple, tractable modeling framework for species coexistence in an ecosystem where habitat heterogeneity is either a fixed feature of the landscape, or dynamically generated through biotic feedbacks. Our models, which are grounded in the metapopulation formalism, add to the rich literature on species coexistence in spatially-varying environments, and advance ecologists’ growing understanding of feedbacks between multispecies communities and environmental variation. For fixed, or exogeneous, heterogeneity, our work extends classic models (Horn & MacArthur 1972; Holt 1997; Shurin *et al*. 2004), recapitulates foundational theory in a new setting (Chesson 1985; Iwasa & Roughgarden 1986; Chesson 2000), and formalizes a connection between habitat partitioning and consumer-resource dynamics. In the context of endogenous heterogeneity, our model demonstrates how environmental modification by different species can lead to the stable maintenance of landscape and species diversity over long time scales, and clarifies the conditions under which these processes will occur. The minimal nature of our model makes it a promising tool to understand generic aspects of such feedbacks, which have been studied in many system-specific contexts (Bever *et al*. 1997; Kucharski *et al*. 2016; Ratzke & Gore *2018)*.

Crucially, our approach also allows us to study exogenous and endogenous heterogeneity in a shared framework, revealing similarities and differences between coexistence mediated by both kinds of environmental variation. We showed that coexistence under endogenous heterogeneity is sensitive to the distribution of patch types in a landscape, but that the stability of a coexistence equilibrium is insensitive to the structure of species’ colonization and establishment rates. When heterogeneity is endogenously generated, in contrast, the ecosystem has the capacity to “self-tune” the distribution of patch states to ensure feasibility. However, the landscape will only evolve toward this coexistence equilibrium if species’ patch modifications generate negative frequency-dependent feedbacks. We derived quantitative criteria for such feedbacks to maintain coexistence, and showed that these same criteria apply whether species modify the environment on very short or very long timescales. While the model dynamics are not identical at these two extremes, there are qualitative similarities that distinguish the endogenous cases from the exogenous one.

Although our models are quite abstract, they may nonetheless be useful for inference in natural systems. Eq. 4 expresses a relationship between the joint distribution of patch types and resident species at equilibrium (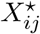), and the model parameters *p*_*ij*_. An identical relationship holds for Eq. 5 once the distribution of patch states has equilibrated. We have discussed how this relationship expresses a net pattern of habitat association that emerges from the combined effects of species sorting and mass effects. By inverting this relationship, one can also obtain an estimator for each *p*_*ij*_ parameter (up to a constant factor, *m*) in terms of 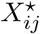 and the marginal frequencies 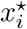 and 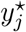. In principle, these frequencies could be computed in systems where well-resolved census and environmental data are available. Such data have been collected for plant communities (Harms *et al*. 2001; Phillips *et al*. 2003). This method makes it possible to estimate dynamical model parameters from static, observational data, which is highly desirable for systems such as tropical forests where the slow pace of dynamics complicates experimental or time-series analysis. However, an important challenge to putting this approach into practice is defining operational patch types in context.

Our results suggest other practical considerations for inference and management in natural landscapes. For example, we found that manipulating the distribution of patch types in a landscape (with exogenous heterogeneity) can produce unexpected changes in the distribution of species frequencies. In particular, we illustrated a case where increasing habitat diversity quantitatively decreased species diversity qualitatively, even though all species were specialized on different habitat types (Horn & MacArthur 1972; Shurin *et al*. 2004). Without a very accurate knowledge of *P*, it is difficult to predict the effects of a perturbation to the landscape, and without considering the entire community context, it is impossible to determine important practical thresholds, such as the minimum abundance of a habitat type needed to sustain a species that relies on it. As another example, we showed that biotic feedbacks can produce situations where all species are transiently resilient to perturbations, despite longterm dynamics that drive some of them to extinction. In such systems, efforts to characterize stability using experimental perturbations or analysis of population fluctuations over short timescales may be misleading about stability over longer timescales (Hastings *et al*. 2018).

Of course, the abstract formulation of our model means that significant caveats may apply. We sought to explore the relationships between habitat heterogeneity, environmental feedbacks, and species coexistence in a minimal framework for these processes; therefore our model necessarily neglects other important processes present in real ecosystems. Following other patch models, our framework assumes no local co-occurrence of species, and therefore no direct species interactions. This allows us to isolate dynamics mediated by landscape heterogeneity, but direct interactions are certainly crucial in many systems, and within-patch dynamics might interact with regional dynamics in non-trivial ways. We also treat space implicitly in our model, but the spatial structure of ecological landscapes could shape how patch modification and dispersal interact. For example, a recent theoretical analysis incorporating both habitat partitioning and Janzen-Connell effects (endogenous heterogeneity) showed that these processes can combine to promote coexistence in a strongly synergistic manner when the spatial autocorrelation of patches is accounted for (Smith 2022). Even when only one source of heterogeneity is present, the spatial arrangement of patches could modulate our understanding of coexistence, for instance by reducing source-sink effects that limit the coexistence of imperfect habitat specialists (Snyder & Chesson 2003).

Our approach also relies on a highly idealized implementation of environmental modification and legacy effects. We assume that patches can be classified into discrete types, and when heterogeneity is generated by the community, we assume that each species’ effects on the environment can be summarized by a single patch state. This discreteness will only approximate the variation of environmental conditions in some natural systems, although in others it may be apt. For instance, plant-associated microbial communities can differentiate into discrete types (Bever 2002; Schweitzer *et al*. 2008; Philippot *et al*. 2013). We also assume that when environmental conditions change, they do so by shifting abruptly between these discrete states. In reality, local conditions may change gradually, potentially reflecting the legacy effects of multiple past resident species at a single time, especially when change is slow (e.g., in the *d* → 0 limit).

Intriguingly, one case where these simplifying assumptions might apply directly even in the *d* → 0 limit comes from multi-strain pathogen systems (Kucharski *et al*. 2016). It has long been known that a host’s first infection by a pathogen can induce a lifelong imprinting effect, specific to the strain of first infection, that shapes future immune response (Oidtman *et al*. 2021). This phenomenon, known as original antigenic sin, can have consequences at the level of population and strain dynamics. Recasting individual hosts as patches, and imprinting effects that modulate susceptibility as patch states, our model maps naturally onto these dynamics. Here, changes in the “landscape” occur not through shifts between states, but through demographic turnover that replaces imprinted hosts (lost to mortality) with naive, newborn hosts on timescales much longer than infection dynamics. In SI Appendix 2.4, we show that an alternative model incorporating these processes behaves qualitatively identically to the model studied above. Our modeling approach could help explain how imprinting effects affect the maintenance of strain diversity. Interestingly, (Miller & Allesina 2021) showed that our stability criterion for symmetric feedbacks would imply a “burden of diversity” in such systems – greater strain diversity would raise the threshold for interventions aimed at eradicating the pathogen, a phenomenon that has been investigated in some endemic pathogen systems (He *et al*. 2021).

This example aside, our mathematical description of environmental modification is likely to be just a coarse approximation in many systems. Still, our approach provides a tractable way to link environmental conditions and community composition across varied timescales, offering a first step toward a more complete picture of these landscape feedbacks in a multispecies context. Unlike most existing approaches, our model can provide analytical guidance for such dynamics, and is even fully solvable in special cases. It has become increasingly clear that environmental feedbacks play a ubiquitous role in mediating interspecific interactions, whether maintaining species coexistence (Bever *et al*. 1997; Wright *et al*. 2002), shaping patterns of abundandance and productivity (Roughgarden 1978; Mangan *et al*. 2010; Monk & Schmitz 2021), or driving species invasions (Wolfe & Klironomos 2005; Van der Putten *et al*. 2013; Amor *et al*. 2020). Ecosystems are rarely a one-way street from landscape variation – set by abiotic features or “bottom-up” biotic process – to species diversity; understanding the dynamic interplay between environmental heterogeneity and “top-down” habitat modification is an important goal for ecology (Crooks 2002; Pastor 2005; Monk & Schmitz 2021). Simple models for these processes can illuminate essential features of each source of heterogeneity, and guide the way to unraveling how they interact and contribute to structuring natural communities.

### Materials and methods

All computations were conducted in R (version 3.6.3). Where dynamics are shown, the model equations were integrated numerically using the deSolve package with the ode45 RungeKutta method. Code to generate all figures and simulation results, including all parameters used in the text, is available on GitHub (Miller & Allesina 2022).

## 5 Supporting information

These supplementary materials are organized by the type of heterogeneity. We provide derivations, analysis, and commentary for the results stated in the Main Text.

### 5.1 Exogenous heterogeneity

In these sections, we derive and analyze the mathematical models for exogenous environmental heterogeneity (Eqs. 3.1-3.2) that appear in the Main Text.

#### 5.1.1 Model reduction

We begin with the mathematical model

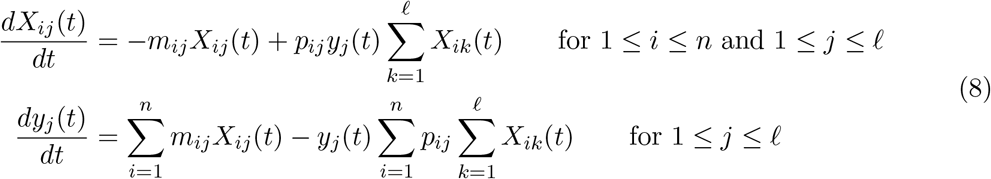

which is introduced and motivated in the Main Text. These dynamics can be viewed as a straightforward extension of classic metapopulation models to include *ℓ* patch types and *n* species, with colonization rates *p*_*ij*_ that depend on the combination of patch type and species identity.

The total number of patches of each type, whether vacant or occupied, is constant through the dynamics. Defining these quantities by *w*_*j*_ = *y*_*j*_(*t*) + ∑_*i*_ *X*_*ij*_(*t*), we have that 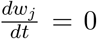, which can be verified by summing the derivatives in Eq. 8. Naturally, then, we also have ∑_*j*_ *w*_*j*_ constant, and we take this total to be 1, so that all model variables can be interpreted as proportions, or frequencies.

Assuming *m*_*ij*_ = *m*, as in the Main Text, we can simplify Eq. 8 by summing over patch types in the *X*_*ij*_. We find that

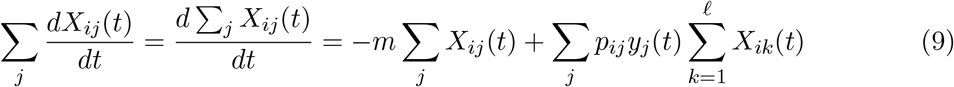

and, defining *x*_*i*_(*t*) = ∑_*j*_ *X*_*ij*_(*t*), the total proportion of patches occupied by species *i*, we have the reduced model

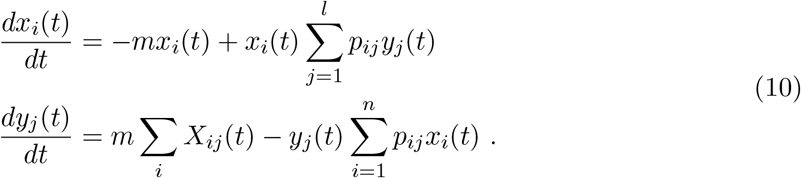

Eq. 10 still contains the quantities Σ_*i*_ *X*_*ij*_(*t*), which are the total proportions of occupied patches of type *j*. Because the total proportion of patches of each type is constant, we can express Σ_*i*_ *X*_*ij*_(*t*) = *w*_*j*_ − *y*_*j*_(*t*) and re-write Eq. 10 only in terms of *x*_*i*_(*t*) and *y*_*j*_(*t*) variables:

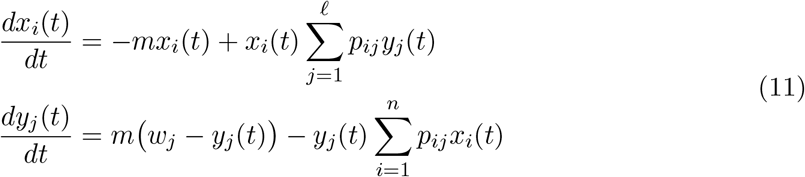

This is Eq. 2 in the Main Text. Notice that Eq. 11 does not uniquely determine the dynamics of Eq. 8. However, we will see that a stable equilibrium for Eq. 11 implies a stable equilibrium solution for Eq. 8, so we concentrate on studying Eq. 11. As noted in the Main Text, Eq. 11 is typically a much smaller system than the full dynamics, because the size of this model grows with the sum of the number of patch types and species, rather than the product. Eq. 11 is also formally equivalent to widely-studied consumer-resource models with abiotic resources, making it possible to leverage known properties of such models.

#### 5.1.2 Coexistence equilibrium and feasibility

In order to study the coexistence equilibrium (i.e. equilibrium where all species are present at positive abundances) of Eq. 11, it is convenient to express the model in matrix form. Using the notational conventions of the Main Text, we have

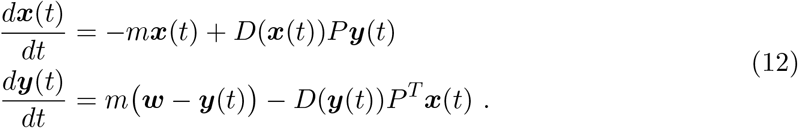

Here *P* is an *n* ×*𝓁* matrix. As discussed in the Main Text, coexistence of all species requires that *n* ≤ *𝓁*. We focus on the case where *n* =𝓁, and so *P* is a square matrix.

However, we note that there is a growing literature on typical properties of this model (in the consumer-resource context) when *n < 𝓁*, which could be applied to understand coexistence in heterogeneous landscapes using the equivalence established here (Tikhonov & Monasson 2017; Cui *et al*. 2020). We will also assume that *P* is invertible, which requires that there are differences among species in their ability to colonize different patch types.

Under these assumptions, the coexistence equilibrium frequencies can be obtained by setting the derivatives in Eq. 12 to zero, and then solving for ***x***^⋆^ and ***y***^⋆^. We find

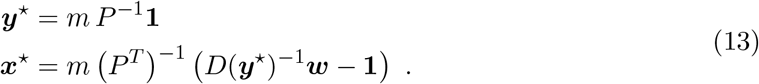

Feasibility of this equilibrium requires that all *P* ^*−*1^**1** *>* 0, and that 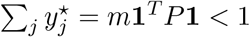 The latter condition effectively sets an upper bound on the local extinction rate, *m*; the condition can always be met if this rate is sufficiently small. Feasibility of the ***x***^⋆^ is also sensitive to the distribution of patch types, ***w***, as explored in the Main Text. Our analysis of the model with endogeneous heterogeneity, below, will imply that, if ***y***^⋆^ is feasible, there is always at least one admissible (i.e. positive and summing to one) ***w*** such that ***x***^⋆^ is feasible. Thus, the condition *P* ^*−*1^**1** *>* 0 plays a fundamental role in determining whether species coexist.

We briefly consider the interpretation of this condition in ecological terms. Fully characterizing the relationship between *P* and feasibility is a difficult problem with no simple account despite significant study of the same underlying mathematical relationship in other contexts (Saavedra *et al*. 2017; Serván *et al*. 2018; Saavedra & AlAdwani 2021; Pettersson *et al*. 2020). This condition expresses the fact that *P* ***y*** = **1** must have a positive solution, or in other words, there must be some possible distribution of available patch types such that all species attain equal effective colonization rates (i.e. colonization rate averaged over the distribution of available patch types: ∑_*j*_ *p*_*ij*_*y*_*j*_). More loosely, we can interpret this condition as requiring that all species are sufficiently similar in overall colonization ability, or sufficiently specialized on distinct habitat types, or some combination of the two.

To justify this statement, we first re-write *P* in a more interpretable form. One can think of 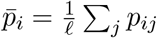 as summarizing the overall colonization ability of species *i* – this is the average colonization rate for species *i* across all patch types. We can express *P* as

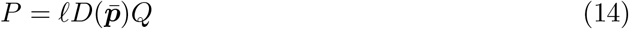

where *Q* is a row stochastic matrix. The elements of *Q* represent the *relative establishment ability* of each species in different patch types. Intuitively, *Q* captures the degree of specialization of each species – if the elements of *Q* are of similar magnitude within a row, the corresponding species is not strongly specialized on a particular habitat.

Now we can show that if all species are exactly equal in overall colonization ability, or perfectly specialized on distinct habitat types, then coexistence is always feasible. In the first case, 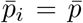 for all species, so 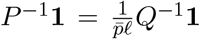. Because *Q* is row stochastic, *Q*^*−*1^ is also, and so 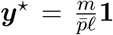, which is all positive. In the second case, *Q* is the identity matrix, so *P* is a positive diagonal matrix, and 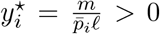. These two extreme cases show that when all species have equal overall colonization ability, coexistence is feasible regardless of specialization, and similarly when all species are perfectly specialized, coexistence is feasible regardless of overall colonization abilities.

Next we develop a quantitative version of the statement that some combination of equality and specialization is necessary for feasibility. Assume that all species are able to colonize and establish in all habitat types at some non-zero rate. Let *q*′ be the minimum element of *Q* (i.e. the smallest relative establishment ability of any species in any patch type). *q*′ provides one measure of specialization in the community. Larger *q*′ implies less overall specialization, while small *q*′ means that there is at least one species that relies on certain habitat types much less than others. We prove that there is now a limit on how much any two species can differ in their overall colonization ability before feasible coexistence is precluded. Assume that two species, *i* and *j*, have 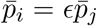. For feasible coexistence, there must be some positive ***y*** such that

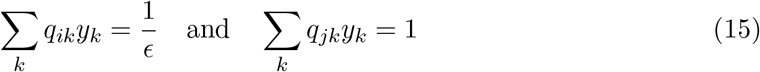

From the former equality we have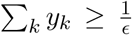, because the elements of *Q* are between zero and one. Then we know that the latter sum is at least equal to 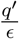. If *ϵ < q*′, then this sum is greater than one, which contradicts the equilibrium conditions. Thus, any two species in the community must be have overall colonization abilities within a factor of 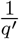, and we have a quantitative statement of the notion that a certain degree of specialization requires a sufficient amount of similarity among species for coexistence. We make no attempt to sharpen this bound; we simply emphasize that these two ecologically meaningful attributes are constrained by one another.

#### 5.1.3 Stability analysis

It is known, in the context of consumer-resource theory, that the coexistence equilibrium of Eq. 11 is always globally stable if it is feasible (see e.g. (Marsland III *et al*. 2020)). Global stability means that trajectories of the system will approach the equilibrium from any initial condition where all species are present. For completeness, we outline a proof of this fact by constructing a Lyapunov function for the dynamics.

A Lyapunov function for an autonomous dynamical system 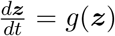, with ***z*** ∈ ℝ^*n*^, is a continuous scalar function *V* : ℝ^*n*^ → ℝ that has continuous first derivatives and satisfies the following:

*1. V* (***z***) *>* 0 except for an equilibrium point ***z***^⋆^, where *V* (***z***^⋆^) = 0

2. 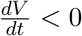 except when ***z*** = ***z***^⋆^, where 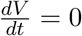

In words, *V* is a positive function that is strictly decreasing through the dynamics, until reaching a minimum at ***z***^⋆^. The existence of such a function implies that the point ***z***^⋆^ is globally Lyapunov stable. For a thorough discussion of Lyapunov functions and global stability, see (Boyce *et al*. 2017; Hahn *et al*. 1963).

Unfortunately, there is no general method to construct Lyapunov functions. However, we illustrate how one can sometimes be found using a technique called *Generalized Lotka-Volterra (GLV) embedding*, which exploits the facts that (i) a large class of differential equations can be recast as GLV systems, and (Szederkenyi *et al*. 2018) (ii) well-studied candidate Lyapunov functions for GLV systems are known (Goh 1977). Thus, GLV embedding removes some of the need for “divine inspiration” that is usually require to find Lyapunov functions. For more detailed exposition of this method, see (Rocha Filho *et al*. 2005; Szederkenyi *et al*. 2018; Miller & Allesina 2021).

Eq. 11 is not a GLV system, but we can define the new variables 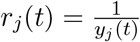 and “recast” the model as the larger system

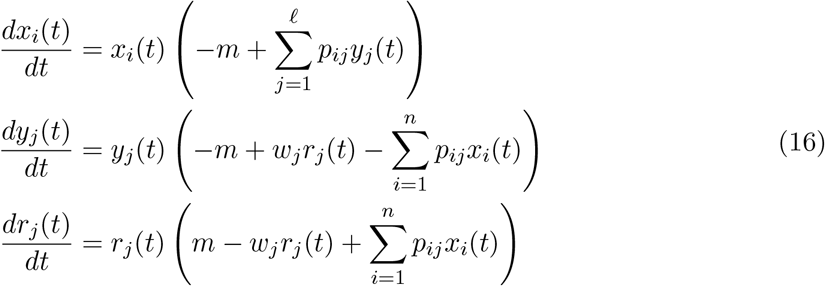

subject to the constraints 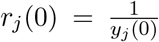 on the initial conditions. This system *is* a GLV model. If we define the vector of abundances ***z***(*t*) = (***x***(*t*), ***y***(*t*), ***r***(*t*)), Eq. 16 can be written as 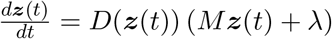 with

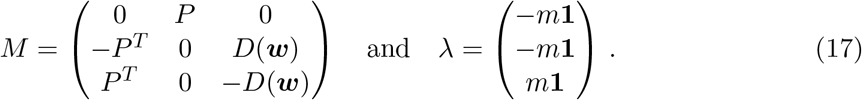

Now we that we have a GLV system, we can try to apply the well-known candidate Lyapunov function (Goh 1977)

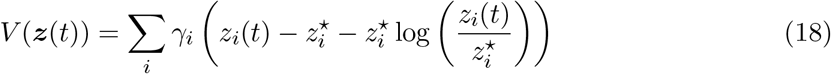

where ***γ*** is a vector of non-negative constants, and 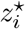 is the equilibrium value of the corresponding component of ***z***. Defined this way, *V* is positive everywhere except at the coexistence equilibrium (Goh 1977). The time derivative of *V* is given by

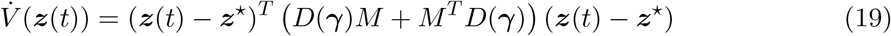

and thus we seek a choice of ***γ*** such that 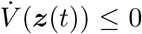 for all times. Consider *γ* = (**1, 1, 0**)^*T*^. Then we find

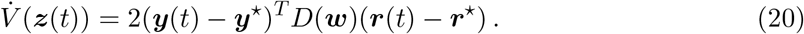

But we know that 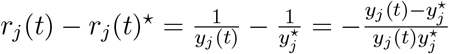 and so

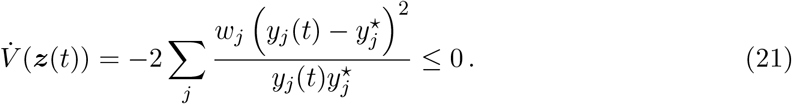

This expression is actually zero whenever ***y***(*t*) = ***y***^⋆^, regardless of whether ***x***(*t*) is also at equilibrium. But we note that if ***y***(*t*) = ***y***^⋆^ and ***x***(*t*) ≠ ***x***^⋆^, then 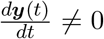 and the trajectory leaves the set where 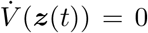. Thus, LaSalle’s invariance principle guarantees that the coexistence equilibrium is globally stable.

The stability of this equilibrium for Eq. 11 immediately implies that the full model in Eq. 8 also reaches a stable coexistence equilibrium. Taking ***x*** and ***y*** at equilibrium, the dynamics for the joint distribution of species and patch types become:

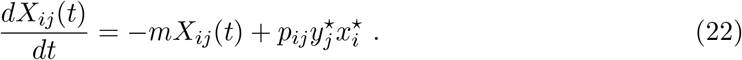

These equations are now uncoupled from one another, and each has a stable (noting that –*m* is always negative) equilibrium given by Eq. 4 in the Main Text.

### 5.2 Endogenous heterogeneity

In these sections, we derive and analyze the mathematical models for endogenous environmental heterogeneity (Eqs. 3.5-3.7) that appear in the Main Text. We also introduce an alternative model where environmental modification is tied to demographic turnover in a host population, or strong disturbances that revert patches to a “naïve” state. We show that our major conclusions are robust to this difference in model particulars.

#### 5.2.1 Model reduction

As described in the Main Text, we now consider a scenario where the type of each patch is not fixed, but instead patches can be in one of *n* states, corresponding to the *n* species in the community. When a patch in state *j* is occupied by species *i*, at some rate it may shift to state *i*. This provides a phenomenological way to model habitat modification by each species, which can change the state of the environment over some characteristic timescale. Combining these new processes with the assumptions outlined for dynamics with exogenous heterogeneity, we have a model analogous to Eq. 8, which in the most general case is given by

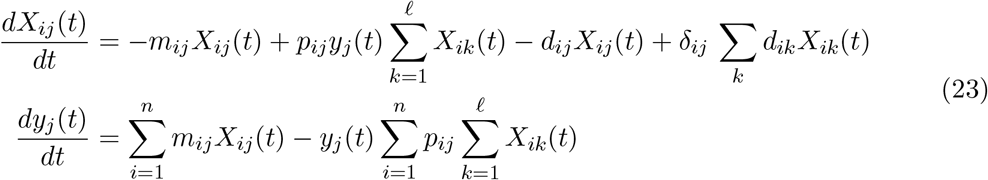

Here *δ*_*ij*_ is the Kronecker delta, which equals one when *i* = *j* and zero otherwise. In this model, we automatically have 1 ≤ *i, j* ≤ *n* by assumption. In general, the rate at which species *i* modifies patches in state *j* might depend on both *i* and *j*, as shown here. Following the exogeneous case, however, we assume *m*_*ij*_ = *m* and *d*_*ij*_ = *d* and track the dynamics for *x*_*i*_(*t*) =∑ _*j*_ *X*_*ij*_(*t*). In this case, we must also track the dynamics of patch states, since these are not fixed. We define *w*_*k*_(*t*) = *y*_*k*_(*t*) +∑_*i*_ *X*_*ik*_(*t*). Using these definitions to compute sums of the derivatives in Eq. 23, we write the reduced model

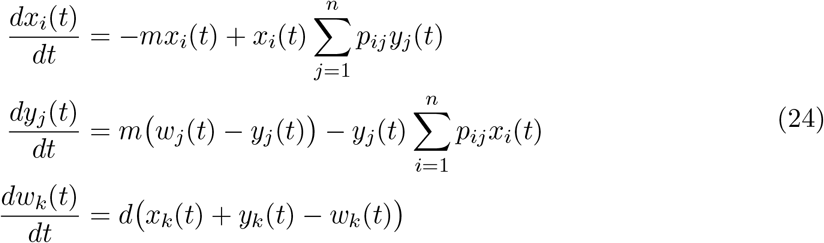

which is Eq. 3.5 in the Main Text.

As described in the Main Text, we can study this model in two limits (*d* → ∞ and *d* → 0, corresponding to very fast and very slow feedbacks) where the equations become simplified. These fast-slow decompositions rely on Tikhonov’s theorem (Tikhonov 1952), and are valid whenever the coexistence equilibrium for the relevant fast system is feasible and stable. We know that stability will hold in all cases, and in the limit *d* → ∞, feasibility is also guaranteed. In this fast limit, Eq. 24 becomes identical to the model studied by (Miller & Allesina 2021). We refer the reader to this study for detailed analysis of the model dynamics. However, we note that Miller and Allesina showed that when *P* = *P*^*T*^, the coexistence equilibrium is stable if and only if *P* has exactly one positive eigenvalue. This condition reappears in the analysis below. Miller and Allesina additionally discuss the ecological interpretation of this eigenvalue condition.

#### 5.2.2 Stability for slow feedbacks

In the slow limit (*d* → 0), we take

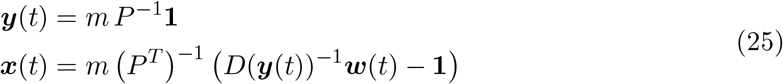

assuming that these fast variables rapidly approach their equilibrium given ***w***(*t*), and then substitute into the equations for 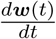 to obtain the slow system

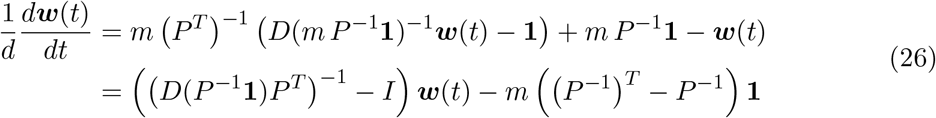

which is Eq. 3.7 in the Main Text. This decomposition holds when ***y***(*t*) and ***x***(*t*) are feasible, which may not be true for all initial conditions ***w*** (see discussion in Main Text).

Eq. 26 is a linear matrix equation with several special properties. The constraint Σ_*k*_ *w*_*k*_(*t*) = 1 implies that the dynamics should be confined to the simplex. These dynamics can be expressed in terms of the eigenvectors and eigenvalues of *A* = (*D*(*P* ^*−*1^**1**)*P*^*T*^)^−1^ − *I*. We note that *D*(*P* ^*−*1^**1**)*P*^*T*^ is column stochastic, so this matrix has a right eigenvector, which we denote ***v***, with an associated eigenvalue of 1. Because the matrix is non-negative, ***v*** is in fact the Perron eigenvector of *D*(*P* ^*−*1^**1**)*P*^*T*^, and it is also non-negative (Horn & Johnson 2012). It is easy to see that *A****v*** = 0, which means that the dynamics have no component in this direction. The remaining eigenvectors of *A* have components that sum to zero, as we verify by considering

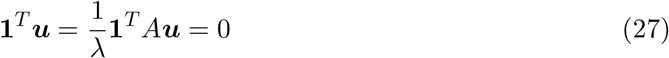

for any eigenvector *u* with associated eigenvalue *λ* ≠ 0. Here we used the fact that **1** is a left eigenvector of *A* with an associated eigenvalue of zero. Because the dynamics can be written as a linear combination of zero-sum eigenvalues, the dynamics for ***w*** are zero-sum, and remain in the simplex, as expected.

Taking this constraint into account, Eq. 7 has a unique equilibrium given by

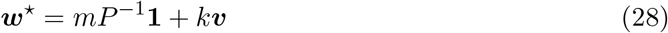

where ***v*** is the eigenvector defined above, normalized such that **1**^*T*^ ***v*** = 1, and *k* is the quantity 1 − *m***1**^*T*^ *P* ^*−*1^**1**. We can verify that

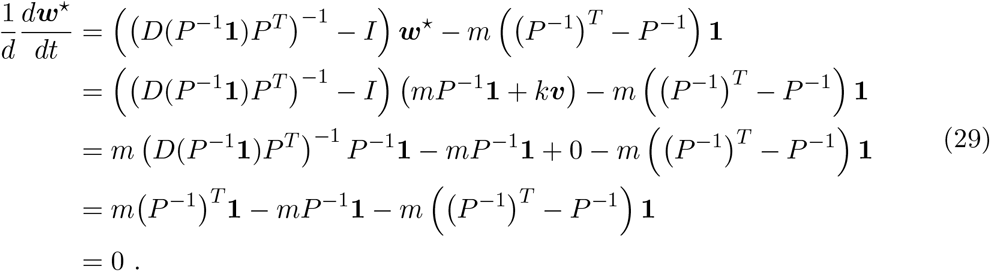

Provided that *P* ^*−*1^**1** *>* 0 and *m***1**^*T*^ *P* ^*−*1^**1** *<* 1 (the feasibility conditions for ***y*** in the fast system), this equilibrium is feasible. It also implies a feasible equilibrium for the species frequencies, given by

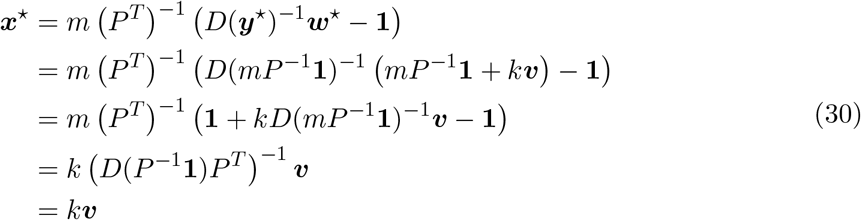

Here we used the definition of ***v***, and the fact that ***v*** is a Perron eigenvector implies that all components are positive.

It is worth noting that because the existence of this equilibrium is independent of the timescales of the dynamics, these equilibrium calculations apply for Eq. 24, for any choice of *d*.

Finally, we consider the stability of this equilibrium (assuming again that *d* →0). Trajectories of Eq. 26 will approach ***w***^⋆^ from any initial condition if and only if the matrix *A* has eigenvalues with all negative real parts (excepting the zero eigenvalue, which corresponds to directions forbidden by the zero-sum constraint). Equivalently, (*D*(*P* ^*−*1^**1**)*P*^*T*^)^*−*1^ must have eigenvalues with real part less than 1. To relate this condition to the eigenvalues of *D*(*P* ^*−*1^**1**)*P*^*T*^ (or, equivalently, *PD*(*P* ^*−*1^**1**), which shares the same eigenvalues), suppose that this matrix has an eigenvalue *a*+*bi*. The eigenvalues of an inverse matrix are the inverses of the eigenvalues, so (*D*(*P* ^*−*1^**1**)*P*^*T*^)^*−*1^ has a corresponding eigenvalue with real part 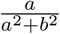. Thus, we require that 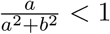 or equivalently 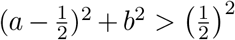. Overall, the Perron-Frobenius theorem implies that *a*^2^ + *b*^2^ *<* 1 (because *D*(*P* ^*−*1^**1**)*P*^*T*^ has an eigenvalue 1), and stability requires that 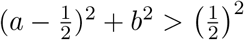. In graphical terms, the eigenvalues of *D*(*P* ^*−*1^**1**)*P*^*T*^ must lie within a crescent-shaped region in the complex plane, within a circle of radius 1 centered at the origin, and outside of a circle of radius 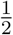 centered at 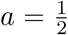 (see Fig. 3.6). From this geometric picture, we immediately see that if all but one eigenvalue (the Perron eigenvalue) of *D*(*P* ^*−*1^**1**)*P*^*T*^ have negative real part, this is sufficient to ensure stability.

**Figure 6:**
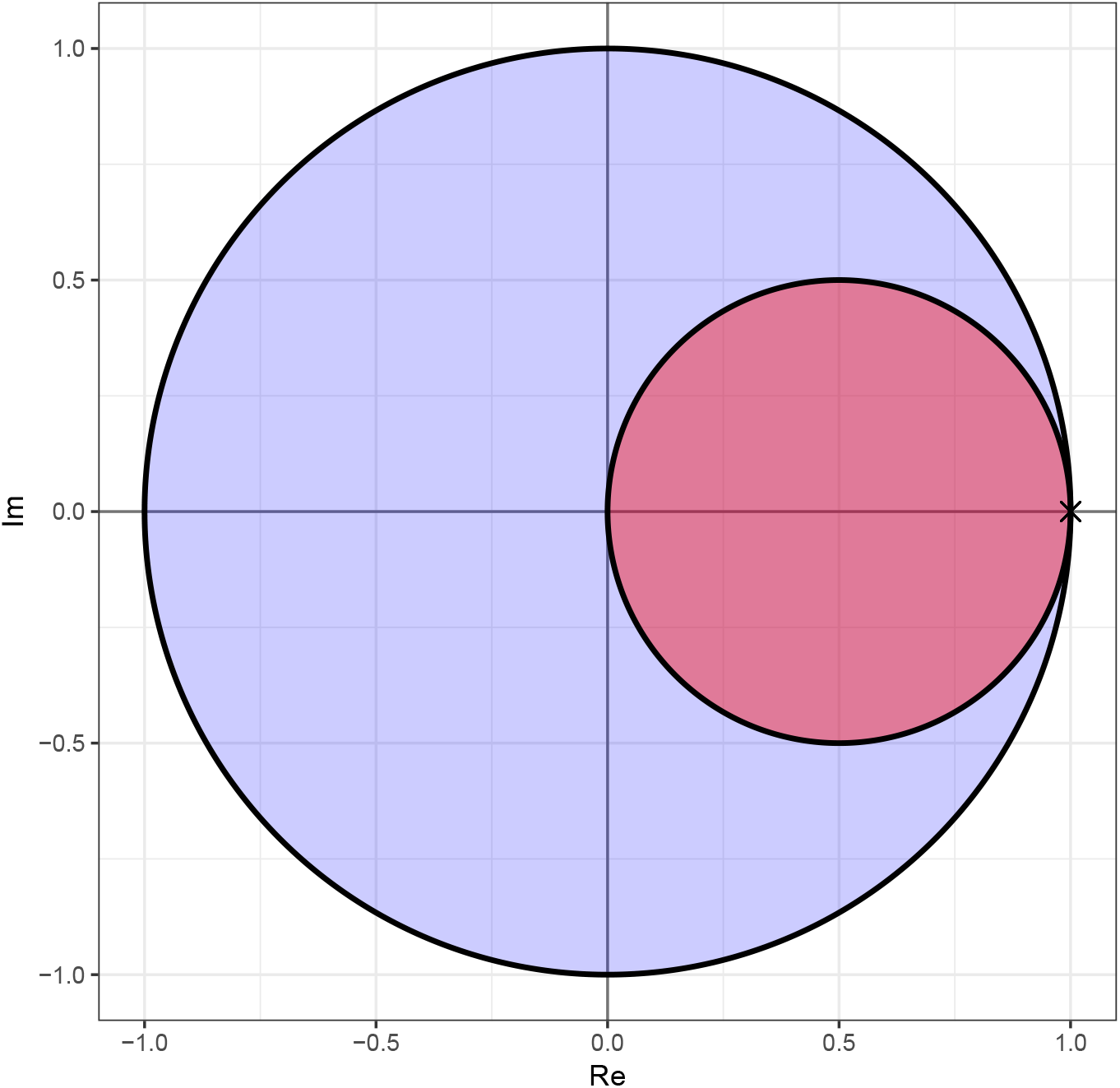
Graphical stability condition for Eq. 26. Stability of the coexistence equilibrium is determined by the eigenvalues of *D*(*P* ^*−*1^**1**)*P*^*T*^ (or, equivalently, *PD*(*P* ^*−*1^**1**)). This matrix has a Perron (dominant) eigenvalue *λ* = 1, indicated here by a black X. The remaining *n* – 1 eigenvalues fall within the unit disk in the complex plane (light blue shaded region). The coexistence equilibrium is stable if these remaining eigenvalues all lie outside of a disk of radius 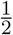 centered at 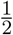 on the real line (red shaded region). Thus, there is a crescent-shaped stability region which must contain the non-Perron eigenvalues in order for the dynamics to be stable. Notice that if all non-Perron eigenvalues have negative real part, then they necessarily lie in the stability region, so this condition is sufficient to ensure stability. When *P* is a symmetric matrix, all eigenvalues of *PD*(*P* ^*−*1^**1**) lie on the real line, so negativity of the non-Perron eigenvalues becomes necessary, as well as sufficient, for stability.

The matrix *PD*(*P* ^*−*1^**1**) defines a natural normalization of *P* in this context – multiplying by the diagonal matrix *D*(*P* ^*−*1^**1**) maps every eigenvalue into the unit disk. In more ecological terms, this normalization also removes any effect of differences in overall quality between patches. To see this, decompose *P* as *QD*(*P*^*T*^ **1**), where *Q* is a column stochastic matrix and *P*^*T*^ **1** are the column sums of *P*. Any differences in the overall quality of patches are captured by *P*^*T*^ **1**. Now we see that *PD*(*P* ^*−*1^**1**) is completely insensitive to *P*^*T*^ **1**: we have *P* ^*−*1^**1** = *D*(*P*^*T*^ **1**)^*−*1^*Q*^*−*1^**1**, and so

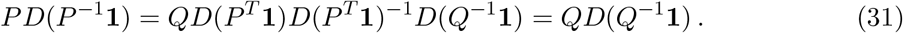

In general, it is not possible to relate this stability condition more directly to *P*. However, in the special case where *P* is symmetric (i.e. *P* = *P*^*T*^), this stability condition reduces to the statement that the coexistence equilibrium is stable if and only if *P* has exactly one positive eigenvalue. This because *PD*(*P* ^*−*1^**1**) is similar to the matrix *S* = *D*(*P* ^*−*1^**1**)^1*/*2^*PD*(*P* ^*−*1^**1**)^1*/*2^ (and therefore shares the same eigenvalues), and *S* is congruent to *P* (and therefore shares the same number of positive, negative, and zero eigenvalues) (Horn & Johnson 2012). This conclusion follows from Sylvester’s law of inertia and relies on the symmetry of *P*. This symmetry also ensures that *P* (and consequently *S* and *PD*(*P* ^*−*1^**1**)) has strictly real eigenvalues. From Fig. 3.6, it is apparent that along the real line, the region of stability reduces to the interval (−1, 0). In other words, the eigenvalues of *P* (excepting the Perron eigenvalue, as usual) must be negative.

#### 5.2.3 Stability for symmetric feedbacks on arbitrary timescales

The appearance of the necessary and sufficient stability condition *P has exactly one positive eigenvalue* for symmetric feedbacks at two opposite limits of the dynamics suggests that this condition should apply more broadly for Eq. 24 with feedbacks on any timescale (i.e any choice of *d*). Here, we prove that this is indeed the case. In particular, we show that this condition characterizes the local stability of the coexistence equilibrium of Eq. 24 whenever *P* is symmetric.

If *P* = *P*^*T*^, the equilibrium frequencies found in the previous section simplify dramatically. We find ***y***^⋆^ = *mP* ^*−*1^, ***x***^⋆^ = *k*^*′*^***y***^⋆^ and ***w*** = (1 + *k*^*′*^)***y***^⋆^, where 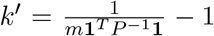. That is, all three equilibrium vectors become proportional to *P* ^*−*1^**1**. This is easily verified by noting that *P* ^*−*1^**1** is a right eigenvector of *D*(*P* ^*−*1^**1**)*P*^*T*^ = *D*(*P* ^*−*1^**1**)*P* with an associated eigenvalue of 1.

This simple form for the coexistence equilibrium of Eq. 24 makes it possible to analyze the Jacobian matrix for these dynamics at equilibrium. We find the Jacobian

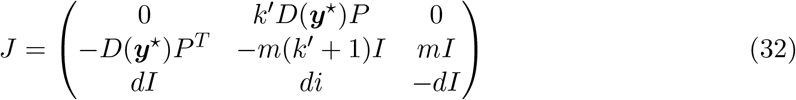

It is actually more convenient to work with another matrix, *J* ^*′*^, that is similar to *J*, and thus shares the same eigenvalues (Horn & Johnson 2012). We first define the change of basis matrix

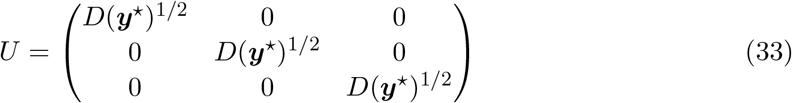

and then define *J* ^*′*^ = *U* ^*−*1^*JU*. More explicitly, we have

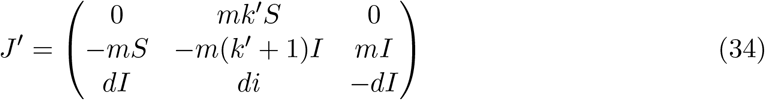

where *S* = *D*(*P* ^*−*1^**1**)^1*/*2^*PD*(*P* ^*−*1^**1**)^1*/*2^. Following (Miller & Allesina 2021), we use the ansatz that the eigenvectors of *J* ^*′*^ all take the form (***u***_*i*_, *c*_1_***u***_*i*_, *c*_2_***u***_*i*_)^*T*^, where ***u***_*i*_ is the *i*th eigenvector of *S*, and *c*_1_ and *c*_2_ are undetermined constants. We will show that each eigenpair of *S* generates three associated eigenpairs of *J* ^*′*^, and the eigenvalues of *S* control the signs of the eigenvalues of *J* ^*′*^.

With our ansatz, each eigenvalue of *J* ^*′*^ satisfies

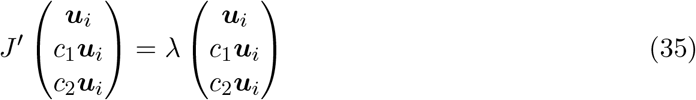

and taking the matrix multiplication by blocks yields the following system of equations:

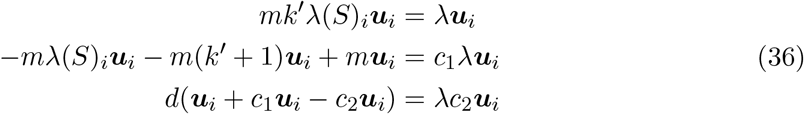

where *λ*(*S*)_*i*_ is the *i*th eigenvalue of *S* (which appears when we consider the definition of ***u***_*i*_). Provided suitable constants exist, this system of equations confirms our ansatz for the form of the eigenvectors of *J* ^*′*^. These equations imply the scalar system

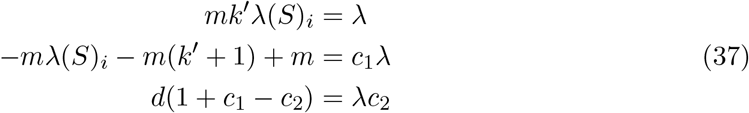

and using substitution for *c*_1_ and *c*_2_, we can reduce this system further to the cubic equation

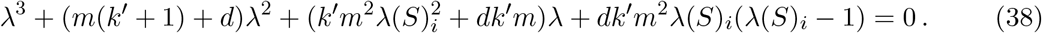

We note that the coefficients of the first three terms are positive for any choice of the model parameters. The constant term is negative if *λ*(*S*)_*i*_ *>* 0, and positive if *λ*(*S*)_*i*_ *<* 0 (noting that the eigenvalues of *S* must be less than 1 in magnitude – see discussion in the previous section). Descartes’s rule of signs implies that number of positive and negative roots is controlled by the sign of the constant terms, and therefore by the sign of *λ*(*S*)_*i*_. If *λ*(*S*)_*i*_ *<* 0, there are zero positive roots; if *λ*(*S*)_*i*_ *<* 0, there is exactly one positive root. There is one special case to consider: we know that *S* has an eigenvalue of 1, and when *λ*(*S*)_*i*_ = 1, the constant term in the equation above vanishes, leading to a root *λ* = 0. This is consistent with our expectation that there is one direction pointing “out” of the simplex, along which the dynamics cannot change.

Overall, this analysis shows that each eigenvalue of *S* generates three eigenvalues of *J* ^*′*^, and the number of positive eigenvalues of *J* ^*′*^ is exactly one less than the number of positive eigenvalues of *S*. In particular, *J* ^*′*^ has all non-positive eigenvalues if and only if *S* has exactly one positive eigenvalue. Because *S* and *P* are symmetric and congruent, Sylvester’s law of inertia dictates that the number of positive eigenvalues of *P* and *S* are the same, and we recover the conjectured stability condition (Horn & Johnson 2012).

#### 5.2.4 An alternative model

In some cases, it is natural to assume that changes in patch state are mediated by significant disturbance events that remove any species present and reset the patch to a “naïve” state. In the context of immune imprinting, discussed in the Main Text, this might actually represent the replacement of an old patch (host) by a new one, due to births and deaths in the host population. In other contexts, we might simply imagine severe perturbations, such as fire, that clear the patch. Then the new state is determined by the next species to occupy this naïve patch.

In this section, we introduce a modification of Eq. 24 that incorporates this kind of process. We show that this modified model behaves qualitatively like Eq. 24 when disturbance events are very rare and naïve patches are re-colonized sufficiently quickly.

We consider the model

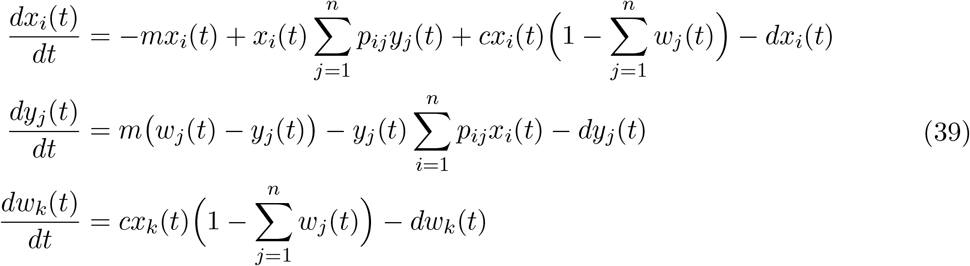

where all variables have the same interpretation as before. Now *d* is interpreted as the rate of disturbance in the system, and the new parameter *c* is the rate at which all species colonize naïve patches. The frequency of naïve patches is represented implicitly as 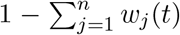, which we henceforth denote by *z*(*t*) for concision. In this model, all patches are subject to disturbance, which leads to losses from the ***x*** and ***y*** variables, and changes in the distribution of patch states (***w***) occur as naïve patches are colonized by different species, which set the state of the patch.

In the limit where *d* → 0 and assuming *z*(*t*) is of the same order as *d*, we can apply a fast-slow decomposition and show that the dynamics of Eq. 39 are essentially identical to the slow limit of Eq. 24. By taking *z*(*t*) to be small, we are assuming that naïve patches are recolonized sufficiently quickly (relative to *d*) so that the frequency of naïve patches in the landscape is small. We will show that this assumption is consistent with the equilibrium behavior for *z*(*t*).

In this limit, we take 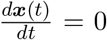 and 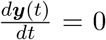, assuming that these variables change and equilibrate much more rapidly than the ***w***(*t*) (more formally, we notice that all terms in 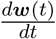 are of order *d*, taking *d* → 0). Then we obtain

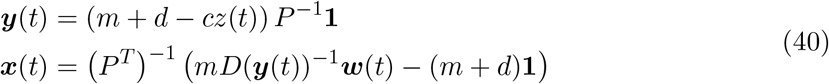

which is nearly identical to equation Eq. 25. Now we consider the dynamics of *z*(*t*), given by

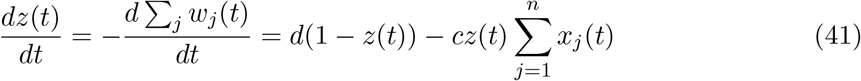

Using Eq. 40, we have

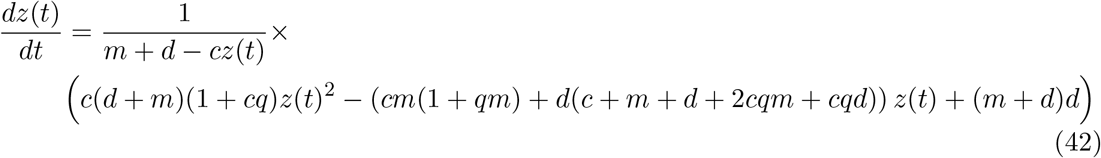

where *q* = **1**^*T*^ *P* ^*−*1^**1**. Under our assumptions that *d* and *z*(*t*) are small, the prefactor is approximately equal to 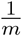, and in particular, it will be positive. This means the dynamics of *z*(*t*) are controlled by the quadratic factor. This factor can be well-approximated by

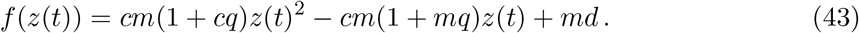

One root of *f* (*z*(*t*)) is small – using a binomial approximation, this root is approximately 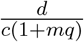. This root is associated with a stable equilibrium point for 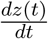, which we can verify by noting that *f* (*z*(*t*)) is negative above this root and positive below. Thus, our initial assumption that *z*(*t*) is of order *d* is supported. For *z*(0) small enough (precisely how small will depend on the other root), *z*(*t*) will approach an equilibrium 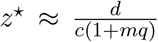. In other words, provided the proportion of patches in the naïve state is initially small, it will remain small through the dynamics.

Finally, we can consider the slow system

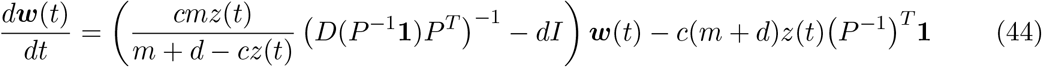

After sufficiently long times, we can take *z*(*t*) ≈ *z*^⋆^ in these dynamics. In this case, Eq. 44 becomes a linear matrix equation, very similar to Eq. 26. The matrix that multiplies ***w***(*t*) shares the same eigenvectors as the matrix *A* in Eq. 26; however the eigenvalues of the two differ. As we have seen previously, (*D*(*P* ^*−*1^**1**)*P*^*T*^)^*−*1^ has an eigenvalue of 1, and so the matrix coefficient in this case has an eigenvalue

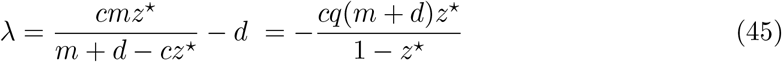

where the last equality follows from the equilibrium condition for *z*(*t*) (i.e. setting 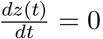). This eigenvalue is always negative, consistent with our conclusion that the proportion of naïve patches, and so the sum of ***w***(*t*), converges to equilibrium. The remaining eigenvectors of (*D*(*P*^−1^ **1**)*P*^*T*^)^−1^ sum to 0, as shown previously. So, as in the original model, the remaining dynamics occur in a manifold where Σ_*j*_*w*_*j*_ is constant. What are the eigenvalues for these directions? Denoting the eigenvalues of (*D*(*P* ^*−*1^**1**)*P*^*T*^)^*−*1^ by *λ*^*′*^, we have

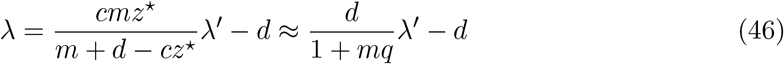

using our approximation 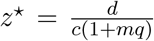. This implies that if Re(*λ*^*′*^) *<* 1 + *mq*, then *λ <* 0. In particular, if Re(*λ*^*′*^) *<* 1, the real part of the corresponding *λ* will always be negative. This is exactly the stability condition we found for the original model, and we see that the conclusions for our original model apply here qualitatively.

